# A Novel Framework to Quantify Dynamic Convergence and Divergence of Overlapping Brain States differentiates Four Psychiatric Disorders

**DOI:** 10.1101/2025.05.20.655164

**Authors:** Najme Soleimani, Sir-Lord Wiafe, Armin Iraji, Godfrey Pearlson, Vince D. Calhoun

**Affiliations:** Tri-institutional Center for Translational Research in Neuroimaging and Data Science (TReNDS), Atlanta, GA; Departments of Psychiatry and Neuroscience, Yale University School of Medicine, New Haven, CT

**Keywords:** Double dynamic functional independent primitive (ddFIP), dynamic functional network connectivity (dFNC), independent component analysis (ICA), mental disorders

## Abstract

Brain function is inherently dynamic, characterized by transient, overlapping functional states rather than static connectivity patterns. Current clustering-based dynamic functional network connectivity methods often fail to capture overlapping states; meanwhile, independent component analysis (ICA)-based methods typically rely on group-level analysis, limiting subject-specific accuracy. To address this gap, we introduce a novel analytical framework estimating individualized dynamic double functional independent primitives (ddFIP)-based states. Our methodological innovation includes: (1) a two-stage ICA combining spatially constrained ICA to define group-level intrinsic connectivity networks (ICNs), followed by constrained ICA to estimate subject-specific states and timecourses; (2) calibration ensuring derived states preserve original correlation scales, enabling meaningful cross-subject and group-level comparisons; and (3) novel metrics leveraging this calibrated representation, including amplitude convergence (uniformity of simultaneous state contributions), amplitude divergence (variability of states independent of state dominance), and dynamic state density (number of concurrently active states at any given time). Validating our framework on an extensive resting-state fMRI dataset (N > 5.5K) spanning four neuropsychiatric conditions revealed disorder-specific connectivity signatures: schizophrenia exhibited extensive variability (increased divergence), while autism displayed pronounced stability (increased convergence). In summary, our proposed method uniquely integrates subject-specific ICA estimation, unit-preserving calibration, and novel convergence-divergence metrics, providing data-driven biomarkers that differentiating psychiatric disorders.

## 1 Introduction

Understanding the human brain as a dynamic, interconnected system involves characterizing how different brain regions communicate and coordinate their activities over time. Functional network connectivity (FNC), derived primarily from functional magnetic resonance imaging (fMRI), quantifies these interactions by measuring correlations among temporal signals of distinct brain regions or networks [Calhoun et al., 2014, Arbabshirani et al., 2013]. Early studies in this domain predominantly utilized static functional network connectivity (sFNC), assuming stable neural interactions averaged across the entire scanning period. However, increasing evidence demonstrates that brain connectivity is inherently dynamic, characterized by continuous fluctuations forming transient and overlapping functional states rather than discrete or constant patterns [Calhoun et al., 2014, Iraji et al., 2021]. This realization has led to the emergence of dynamic functional network connectivity (dFNC) as a promising approach for capturing complex temporal interactions in brain networks, holding significant potential for enhancing our understanding of brain function and dysfunction. dFNC methods capture moment-to-moment variations in brain activity, enabling the identification of temporally distinct connectivity patterns, known as “connectivity states”. These states reflect transient neural interactions among multiple brain networks [Rashid et al., 2014, Dong et al., 2018] and offer a richer and more accurate representation of brain function compared to static measures, thus significantly improving our understanding of neural adaptability and resilience. Accurate characterization of these dynamic connectivity patterns has profound implications for biomarker discovery. Previous methodological approaches in dFNC predominantly rely on clustering-based algorithms, such as k-means, which discretize dynamic interactions into distinct, non-overlapping states. This assumption oversimplifies the continuous and overlapping nature of neural dynamics, potentially missing subtle yet clinically significant patterns, which are critical for capturing true brain dynamics [Rashid et al., 2014, Dong et al., 2018]. Additionally, current independent component analysis (ICA)-based approaches predominantly derive states at the group level Miller et al. [2016], Soleimani et al. [2025], restricting their ability to accurately represent subject-specific variations and preserve original connectivity magnitudes. Consequently, existing methodologies fail to robustly quantify the continuous and overlapping nature of brain connectivity states across subjects and groups.

Dynamic connectivity patterns can particularly be considered as promising biomarkers because they reflect subtle neural variations, allowing for more sensitive detection and characterization of brain disorders [Yan et al., 2024, Soleimani et al., 2025]. Biomarkers are quantifiable biological indicators that can objectively detect physiological and pathological states, facilitating earlier diagnosis, individualized treatment, and precise monitoring of disease progression [Berk, 2023, Le Grand et al., 2022].

Despite substantial prior research investigating the dynamic nature of the brain with application to dFNC-based connectivity biomarkers in various psychiatric conditions, critical methodological challenges persist. For example, studies of schizophrenia (SCZ), bipolar disorder (BPD), major depressive disorder (MDD), and autism spectrum disorder (ASD) often suffer from small sample sizes, limited generalizability, and the inability to capture subtle state overlap and temporal dynamics effectively [Rashid et al., 2014, Nguyen et al., 2017, Shi et al., 2021, Sigar et al., 2023]. Furthermore, existing dynamic connectivity studies frequently rely on predefined networks or limited temporal resolution, restricting their ability to fully realize the potential of dynamic connectivity biomarkers for clinical translation.

To overcome these limitations,we propose an advanced dFNC framework that uniquely integrates both spatially and temporally informed independent component analysis (ICA) to enhance individual-level accuracy. Specifically, we advance existing ICA-based state extraction methodologies through three distinct innovations: (1) Employing a two-stage ICA procedure that first uses spatially constrained ICA to extract intrinsic connectivity networks (ICNs) at the group level, followed by constrained ICA leveraging these ICN templates as priors to estimate accurate subject-specific dFNC states and timecourses, thus preserving individual variability and group correspondence. (2) Implementing a calibration step to ensure the derived subject-specific connectivity states retain the original correlation scales, maintaining interpretability and comparability across individuals and groups. (3) Introducing three novel metrics to comprehensively quantify connectivity state dynamics:

1. *Amplitude convergence* referring to the degree to which multiple connectivity states contribute similarly to the connectivity profile at a given time, indicating a high level of neural flexibility and adaptability.
2. *Amplitude divergence* captures the tendency for connectivity states to contribute at varying levels, reflecting a more uneven distribution of state contributions, which may reflect rigidity or reduced neural adaptability.
3. *Dynamic state density* which quantifies the number of strongly occupied states, reflecting the brain’s preference for spending time in either a smaller or larger subset of dominant states.

These metrics offer additional layers of detail that can characterize different neuropsychiatric conditions based on how flexible or stable their connectivity patterns are over time. By systematically applying and validating the proposed double functional independent primitives (ddFIP) framework on a large resting-state fMRI dataset spanning SCZ, ASD, BPD, and MDD (N > 5,500 participants), we aim to demonstrate the method’s effectiveness in capturing subtle disorder-specific connectivity dynamics and addresses the following gaps explicitly. First, our framework improves methodological rigor by integrating population-derived connectivity templates with subject-specific ICA reconstructions, balancing generalizability and individual variability. Second, it explicitly quantifies continuous fluctuations in connectivity, overcoming limitations imposed by discrete-state clustering methods. Third, by systematically comparing connectivity metrics across multiple neuropsychiatric conditions, our approach facilitates the identification of both disorder-specific and transdiagnostic connectivity patterns.

The significance of this research is twofold. First, from a methodological perspective, this ddFIP-based framework advances our capability to measure dynamic brain interactions accurately, offering the potential for broad application in neuroscientific investigations. Second, clinically, it provides a rigorous foundation for developing reliable dynamic connectivity-based biomarkers, enabling improved diagnostic precision, tailored therapeutic strategies, and deeper understanding of the neural mechanisms underlying psychiatric conditions [Drysdale et al., 2017, Woodward and Cascio, 2015, Yan et al., 2024]. Consequently, this methodological innovation bridges an essential gap between theoretical neuroscience and practical clinical translation, moving toward a biologically-informed and personalized approach to brain research and healthcare.[Zhao et al., 2020, Drysdale et al., 2017, Woodward and Cascio, 2015].

## 2 Materials and Methods

### 2.1 Participants

We assessed our framework using resting-state fMRI data from seven distinct clinical datasets, including the bipolar-schizophrenia network on intermediate phenotypes (B-SNIP) Tamminga et al. [2014], Center For Biomedical Research Excellence (COBRE) Calhoun et al. [2012], Functional Biomedical Informatics Research Network (FBIRN) Keator et al. [2016], a Maryland Psychiatric Research Center (MPRC) study, a Chinese schizophrenia dataset Yan et al. [2019], the Autism Brain Imaging Data Exchange (ABIDE) Di Martino et al. [2014], and MDD dataset collected in Beijing Zhi et al. [2018]. Participants with head motion exceeding 3 ^◦^ or 3 mm were excluded. Additional quality control was performed by assessing the spatial normalization of functional data for each participant, ensuring alignment across the entire brain by comparing individual and group masks. Individual masks were generated by identifying voxels above 90% of the mean brain intensity, while the group mask included voxels present in over 90% of participants. Spatial correlation values were calculated for each subject by comparing the individual mask with the group mask across the top 10, bottom 10, and all slices. Only subjects meeting specific correlation thresholds (0.75 for the top 10 slices, 0.55 for the bottom 10 slices, and 0.8 for the whole brain mask) were retained for analysis, ensuring the inclusion of high-quality fMRI data.

Applying these selection criteria yielded a final sample of N = 5805 participants, diagnosed with SCZ (n=2615; 1302 patients and 1313 healthy controls), ASD (n=1735; 796 patients and 939 healthy controls), BPD with psychosis (n=870; 294 patients and 576 healthy controls), and MDD (n=585; 278 patients and 307 healthy controls). Table 1 provides demographic details of these participants.

**Table 1:** Demographic and Clinical Characteristics of Study Participants

| Disorders | Categories | Age (mean ± std) | Gender (F/M) | Dataset (Sample Size) |
| --- | --- | --- | --- | --- |
| SCZ ($n=2615$) | Patients ($n=1302$) | $35.8 \pm 12.6$ | 487/815 | B-SNIP (429), COBRE (68), FBIRN (151), MPRC (147), 754-sample SCZ (507) |
| | Control ($n=1313$) | $34.2 \pm 11.7$ | 679/634 | B-SNIP (576), COBRE (89), FBIRN (160), MPRC (241), 754-sample SCZ (247) |
| ASD ($n=1735$) | Patients ($n=796$) | $16.7 \pm 9.2$ | 110/686 | ABIDE (796) |
| | Control ($n=939$) | $16.2 \pm 8.6$ | 229/710 | ABIDE (939) |
| Psychotic BPD ($n=870$) | Patients ($n=294$) | $35.4 \pm 12.1$ | 192/102 | B-SNIP (294) |
| | Control ($n=576$) | $36.2 \pm 12.5$ | 341/235 | B-SNIP (576) |
| MDD ($n=585$) | Patients ($n=278$) | $33.4 \pm 11.2$ | 169/109 | 585-sample MDD (278) |
| | Control ($n=307$) | $31.3 \pm 10.4$ | 191/116 | 585-sample MDD (307) |

### 2.2 Neuroimaging Data and Preprocessing

The resting state-fMRI data underwent preprocessing following standard protocols. Using Statistical Parametric Mapping (SPM12; available at http://www.fil.ion.ucl.ac.uk/spm/) within the MATLAB 2022 environment, we initially removed the first ten scans. Motion correction was then applied with a rigid-body transformation tool in SPM to adjust for head movement, followed by slice-timing correction to address acquisition timing differences between slices. The data were subsequently normalized to the Montreal Neurological Institute (MNI) space using an echo planar imaging (EPI) template, with a resampling to 3 × 3 × 3 mm^3^ isotropic voxels. A Gaussian kernel with a full width at half maximum (FWHM) of 6 mm was then applied to smooth the resampled images. Additional processing steps included: (1) detrending to remove linear, quadratic, and cubic trends; (2) de-spiking temporal outliers; (3) Band-pass filtering with a cutoff frequency of 0.15 Hz; and (4) regressing out six head motion parameters and their derivatives. Finally, to reduce potential confounds, we regressed out biological covariates (age and gender) and technical covariates (scanner and site) separately for each disorder.

### 2.3 Spatially Constrained Independent Component Analysis

ICA is a robust, data-driven approach for extracting maximally independent sources from multivariate datasets, widely applied in neuroimaging Calhoun et al. [2009]. Blind ICA, however, encounters the challenge of “order ambiguity,” meaning that the ordering of independent components is arbitrary and can vary between analyses, limiting reproducibility across studies Wang et al. [2014]. While group ICA partially addresses this by establishing correspondence across subjects, it does not fully account for individual variability. An advanced method, spatially constrained ICA, improves upon this by integrating prior spatial information to guide the decomposition, thereby enhancing the precision of IC separation. By applying spatial constraints from anatomical or functional atlases, this technique refines the delineation of brain networks and regions of interest, yielding more interpretable and reliable functional connectivity patterns that are particularly sensitive to subtle changes associated with specific cognitive tasks or clinical conditions Smith et al. [2009], Iraji et al. [2023].

To further ensure consistency in identifying intrinsic connectivity networks (ICNs) across individuals, the NeuroMark pipeline was employed. Using the NeuroMark fMRI 1.0 template Du et al. [2020], as implemented in the GIFT toolbox (http://trendscenter.org/software/gift), 53 ICNs were derived for each scan. The NeuroMark framework leverages spatially constrained ICA to automatically estimate reproducible functional brain markers across subjects and datasets. Unlike region-of-interest (ROI) methods that depend on fixed brain regions, NeuroMark identifies comparable brain networks while accommodating individual variability, which is crucial for assessing group differences and individual classification Tu et al. [2020] Fu et al. [2021]. NeuroMark’s ICNs encompass seven functionally distinct domains: subcortical (SC), auditory (AUD), visual (VIS), sensorimotor (SM), CC, DMN, and cerebellar (CB) components, providing a comprehensive view of brain activity that supports robust inter-subject comparability Du et al. [2020].

### 2.4 Constrained-Dynamic Double Functionally Independent Primitives (c-ddFIP)

A comprehensive workflow of the study in the context of psychiatric disorders using neuroimaging data is provided in Figure 1. This pipeline integrates multiple steps, from data collection to the extraction of dynamic connectivity patterns and the identification of candidate biomarkers. The first step involves collecting and preprocessing fMRI data to extract 53 ICNs from various groups as described elsewhere. These ICN timeseries were utilized to capture dFNC which is an individual 53 × 53 windowed-FNC matrices for each subject using Pearson correlation, a window size of 45s, step size of 1 TR, and a rectangular window. We applied demeaning to each windowed-FNC matrix to remove the static term from the dynamic connectivity signal. This step helps emphasize the temporal variability in connectivity patterns, allowing us to focus on the dynamic changes that occur over time rather than the static baseline connectivity. That is, by removing the static component, we enhance sensitivity to transient fluctuations, which can provide a clearer view of functional connectivity dynamics and improve the detection of subtle connectivity patterns associated with different cognitive states or clinical conditions.

**Figure 1:**
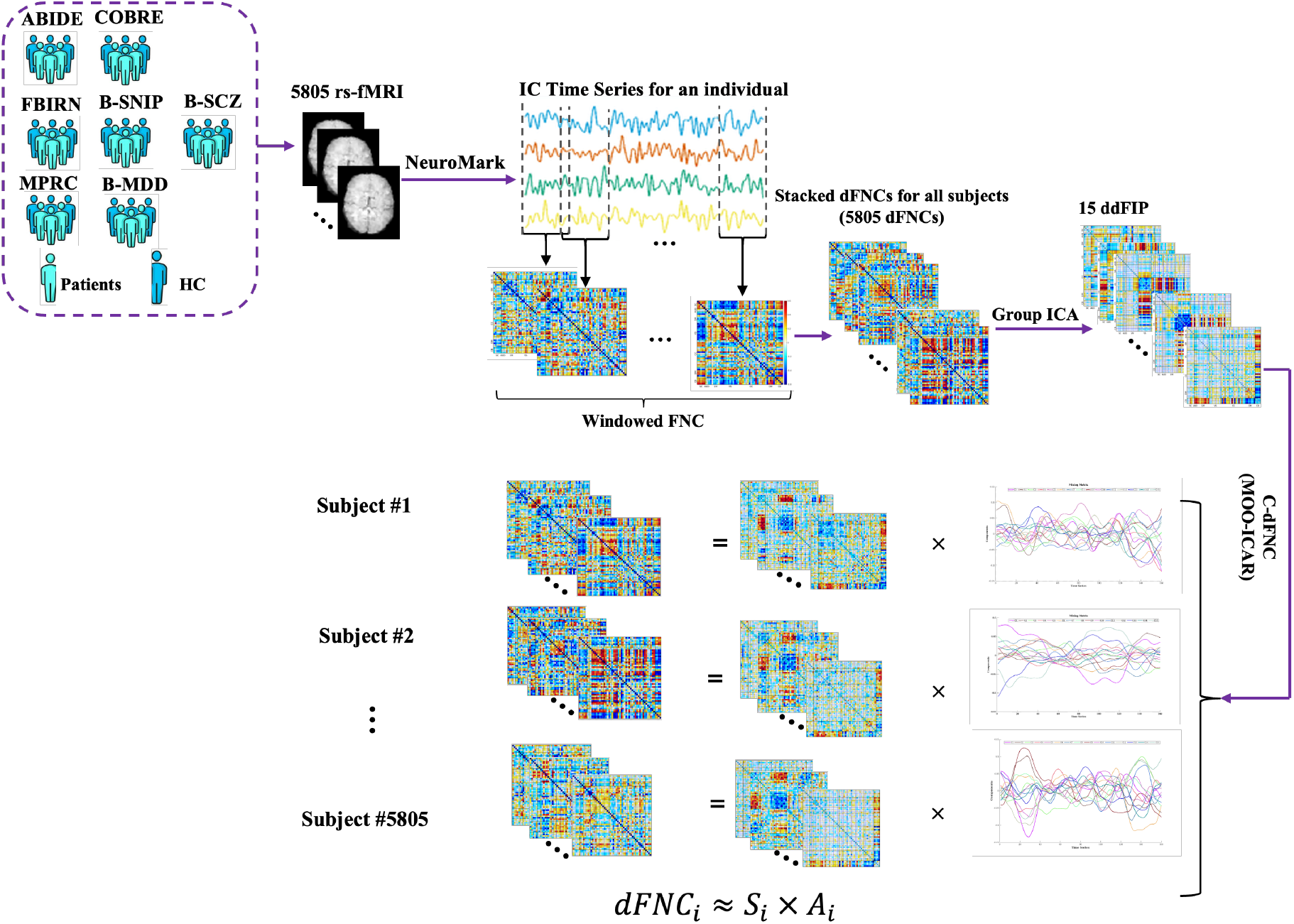
Workflow of the study, rs-fMRI data from both healthy controls and individuals with psychiatric disorders were preprocessed and used to extract 53 ICNs per subject. Dynamic FNC matrices were generated from these ICNs. A blind ICA on concatenated dFNC matrices produced 15 ddFIPs, used as priors in constrained ICA to derive individualized connectivity patterns aligned with population-wide templates.

Individual dFNC matrices are subsequently concatenated across all subjects and time points to form a data set that captures the dynamics of connectivity across the population. Blind ICA is then performed on this concatenated dataset, extracting 15 overlapping states, which function as ddFIPs. These ddFIPs serve as connectivity templates representative of shared patterns across subjects. In other words, these states represent recurring patterns of brain network interactions that can be characteristic of specific psychiatric disorders, which means that the time-varying connectivity profile of each subject is charactereized via a small number of unique states, each corresponding to a particular pattern of brain activity.

We use these 15 ddFIPs as priors in a constrained ICA back-reconstruction process, enabling us to derive individualized connectivity patterns that remain aligned with the population-wide templates. We refer to these constrained source components as constrained dynamic double functional independent primitives (c-ddFIPs). This approach is essential for capturing subject-specific variations in connectivity while ensuring consistency with the broader population dynamics. By applying these shared templates as constraints, we enhance the interpretability of individual results and improve comparability across subjects, which is particularly valuable for detecting subtle differences in connectivity that may relate to specific cognitive or clinical characteristics. Finally, a calibration step is performed to standardize the subject-specific dFNC data by regressing each individual’s dynamic connectivity profile onto predictors formed by the outer product of their mixing matrix and subject-specific states. This yields beta-weighted, variance-scaled timecourses that reflect the contribution of each state, enabling more robust comparisons across individuals and groups while preserving alignment with the population-derived connectivity templates.

### 2.5 Occupancy Analysis

Occupancy analysis provides a measure of how frequently each brain state is most active over time, reflecting the relative stability or prevalence of specific connectivity states within a group. In this study, we computed occupancy by first calculating the dominant state at each time window, defined as the state with the highest relative activity (absolute value) for each individual.

Let *X*_*i*_(*t*) represent the activity of state *i* at time *t*, for *i* = 1, 2, …, *N* states and *t* = 1, 2, …, *T* time windows.

For each time window *t*, the dominant state *C*(*t*) is defined as the state with the maximum relative activity:

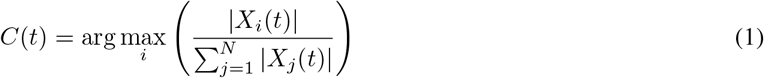

where |*X*_*i*_(*t*) | is the absolute activity of state *i* at time *t*. The normalization by the sum of absolute activities across all states ensures that we assess the relative dominance of each state at each time point.

The occupancy count *O*_*i*_ for each state *i* is the total number of time windows where *i* is the dominant state:

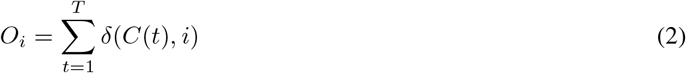

where *δ*(*C*(*t*), *i*) is an indicator function that equals 1 if *C*(*t*) = *i* and 0 otherwise.

The occupancy percentage *P*_*i*_ for state *i* is computed by dividing the occupancy count by the total number of time windows and converting it to a percentage:

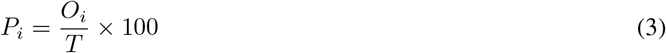

This occupancy percentage *P*_*i*_ represents the proportion of time that state *i* is the dominant activity in the network, providing a measure of its prevalence over time.

By counting the number of time windows during which each state was dominant, we derived the occupancy for each state as the percentage of total time windows. This approach allowed us to quantify the time each state was active, providing insights into the functional connectivity patterns and highlighting the variability and stability of these states across different clinical groups.

### 2.6 Dynamic Convergence/Divergence Framework

In this study, we introduce a novel dynamic convergence/divergence analysis framework designed to assess the stability and variability of brain connectivity patterns over time. This approach is focused on capturing transient shifts in the relative dominance of connectivity states, distinguishing periods when states contribute similarly (converge) or exhibit varying levels of influence (diverge). Unlike traditional static connectivity analysis, which provides a fixed snapshot of connectivity patterns, our framework highlights the temporal dynamics of state amplitudes, offering valuable insights into adaptive or maladaptive neural processes underlying psychiatric disorders such as SCZ, ASD, BPD, and MDD. By focusing on the dynamics of state contributions rather than the stability of connectivity patterns per se, this method moves beyond existing approaches by providing a detailed understanding of the functional flexibility and resilience of the brain in both health and disease.

For each individual, we have a calibrated mixing matrix *X* ∈ ℝ^*T ×N*^ capturing the time-varying activation levels of each state across each window *t*, where *T* represents the number of windows, and *N* represents the number of states (here *N* = 15). To evaluate the dynamic convergence and divergence of these states over time, we define each window at time *t* as

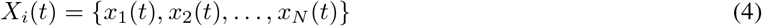

This matrix enables us to track how connectivity patterns evolve dynamically, as each window at time *t* is represented by a feature vector *X*_*i*_(*t*) ∈ ℝ^15^, which contains the weights of the 15 states. To quantify the similarity, or “closeness,” between these states, we calculate the Euclidean distance between any two states *x*_*i*_(*t*) and *x*_*j*_(*t*) at each time point *t* using:

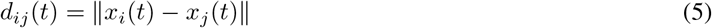

where ∥ · ∥ is the Euclidean norm. This distance metric allows us to rigorously distinguish between moments when brain states have similar amplitudes (converged) or substantially different amplitudes (diverged).

We introduce two thresholds, ϵ and *θ*, to classify states into “converged” or “diverged” categories, respectively. Thresholds are selected based on empirical observations or statistical criteria to ensure they capture meaningful convergence/divergence across subjects and time points. We inspected the distribution of pairwise differences in state contributions across time and subjects to identify natural inflection points indicating low or high divergence. Specifically, states *x*_*i*_(*t*) and *x*_*j*_(*t*) are considered converged if *d*_*ij*_(*t*) ≤ *ϵ*, and diverged if *d*_*ij*_(*t*) ≥ *θ*. We define a function *f* (*d*_*ij*_(*t*)) to classify states *x*_*i*_(*t*) and *x*_*j*_(*t*) into “converged” or “diverged” categories based on the distance *d*_*ij*_(*t*) between them:

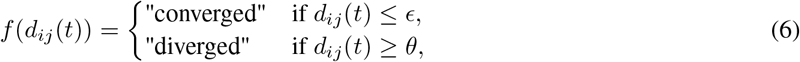

By dynamically tracking these convergence and divergence states, our framework offers an advantage in detecting complex connectivity changes that could reflect cognitive or clinical variations, which may be critical for characterizing mental health conditions.

### 2.7 Dynamic State Density Analysis

We also proposed an additional novel metric focused on the state density. This analysis computes the distribution of “states survived” for different groups based on time-varying connectivity patterns extracted using ICA to identify statistical patterns in different groups. The number of states is denoted as *N*_*c*_ = 15, and the number of time points is *T* = 115. A threshold value (*thr* = 0.06) is applied to define strong states, i.e., states with an absolute value greater than the threshold. For each time point *t* and state *N*, a binary indicator *I*_*t,N*_ is defined as:

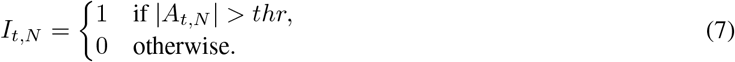

The number of strong states for each time point is computed as:

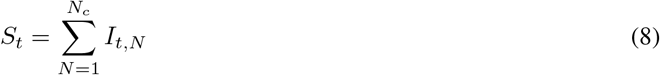

A histogram of *S*_*t*_ is generated using bins 0, 1, …, *N*_*c*_, and the resulting histogram is normalized by the total number of time points (*T*) to yield a probability distribution. The average normalized histogram for each group is computed as:

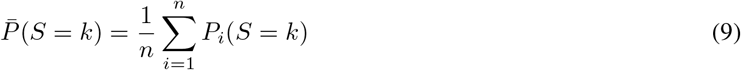

where *n* is the number of subjects in the group, where *P*_*i*_(*S* = *k*) is the normalized histogram for subject *i*, and *k ∈ {*0, 1, …, *N*_*c*_*}* denotes the number of strong states.

To compare patient groups with the HC group, two-sample t-tests are performed for each bin, yielding *p*-values (*p*_*k*_) and *t*-values (*t*_*k*_). The state density analysis captures the distribution of states within dFNC timecourses and identifies significant differences across groups in a dynamic sense, highlighting patterns that differentiate patients with various mental disorders from healthy controls.

We also use k-means clustering of histograms to group similar patterns of state densities across different mental disorders and compare these patterns between patient groups and healthy controls. K-means is an unsupervised machine learning algorithm that partitions the data into a predefined number of clusters, minimizing the variance within each cluster. By clustering the histograms, we can identify distinct patterns or profiles of state densities in the data, which may reflect underlying differences between patient groups and healthy controls, which may help to uncover potential biomarkers or distinctive features associated with different mental disorders.

## 3 Results

### 3.1 Differences in c-ddFIP

To investigate the differences in c-ddFIPs between clinical groups and healthy controls, we conducted two-sample t-tests between patients and healthy controls, across all brain network pairs, represented as cells in our connectivity matrices. Figure 2 provides a comprehensive visualization of cellwise differences between SCZ patients and healthy controls across 15 c-ddFIPs. The results reveal a complex interplay of increased and decreased connectivity across functional networks. Schizophrenia is characterized by hyperconnectivity within default mode and cognitive control networks and hypoconnectivity within sensorimotor and visual networks, highlighting disruptions in sensory integration.

**Figure 2:**
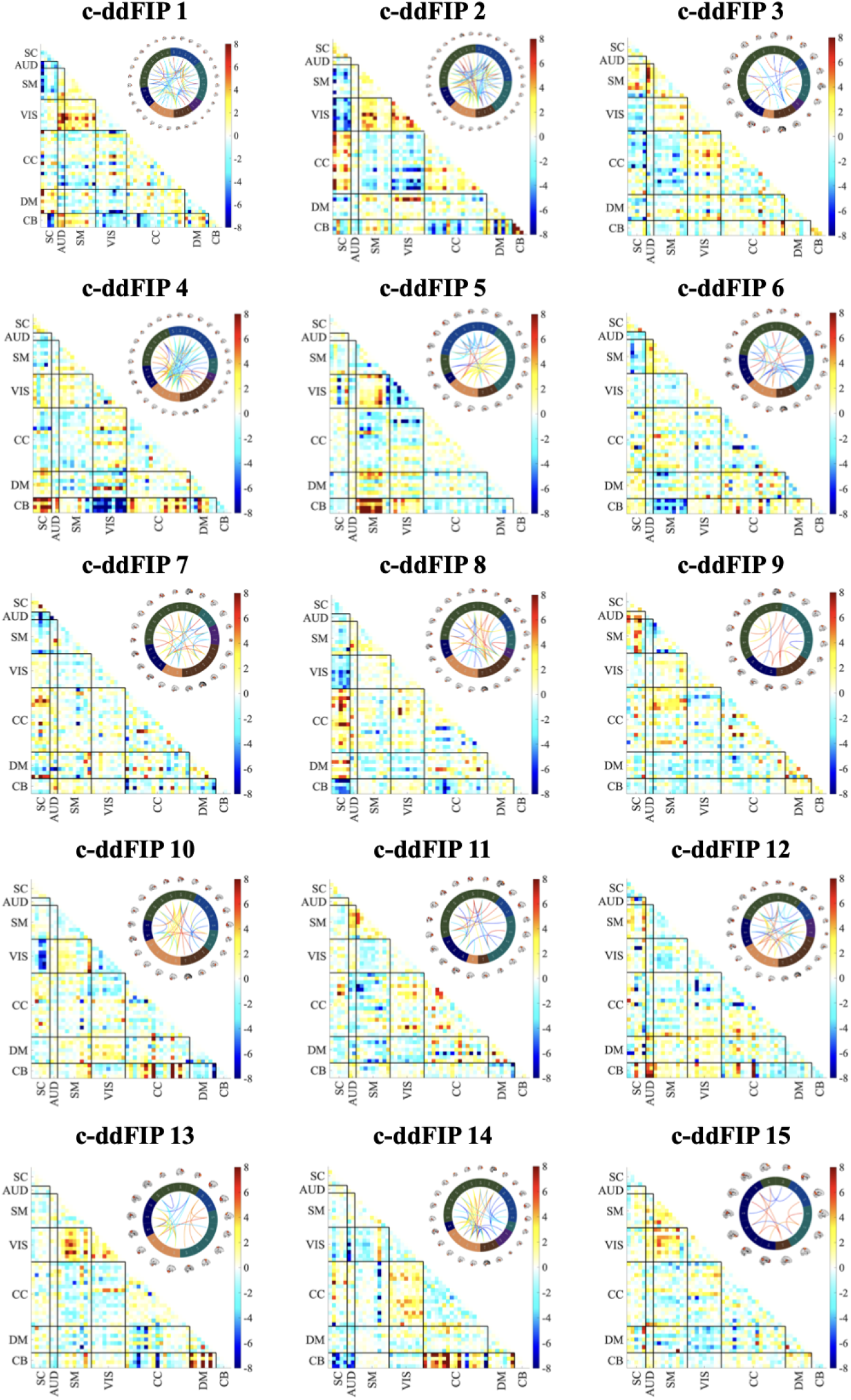
Cellwise differences between SCZ patients and healthy controls (HCs) for each constrained dFIP (c-dFIP), represented as −log_10_(*p*) · sign(*t*). False discovery rate (FDR)-corrected *p*-values (*q <* 0.05) are used to highlight significant differences. Positive values indicate regions with increased connectivity (hyperactivity) in SCZ, while negative values represent decreased connectivity in SCZ. The associated connectograms illustrate the connectivity patterns for each c-dFIP.

However, ASD patients exhibit a dual pattern of hyperconnectivity in higher-order networks within DMN and between DMN and other networks (e.g., CC and AUD), and hypoconnectivity in sensory networks (e.g. SM, AUD, and VIS). Elevated connectivity within and between DMN and CC regions is consistent with symptoms such as persistent negative thoughts and difficulty disengaging from self-focused cognition. SCZ patients show heightened VIS-SM connectivity in c-ddFIP 5, suggesting over-compensatory sensory engagement, while ASD shows reduced VIS-SM connectivity, indicating sensory integration deficits. Furthermore, patients with SCZ exhibit balanced sensory and higher-order engagement, while patients with ASD uniquely emphasize higher-order (DMN-CC) over sensory processing, reflecting a clear divergence. Both BPD and MDD patients exhibit milder disruptions compared to SCZ and ASD.

### 3.2 Differences in Occupancy

The average occupancy percentage plot provides a detailed comparison of how frequently each state is the most active across different clinical groups. The x-axis represents each of the 15 states, while the y-axis indicates the average percentage of time each state was dominant (or “occupied”) for individuals within each group relative to healthy controls. A two-sample t-test performed at the individual level revealed statistically significant differences (*p < 0*.*05*) in specific states across the disorders. These results are consistent with the directional trends seen in Figure 3.

**Figure 3:**
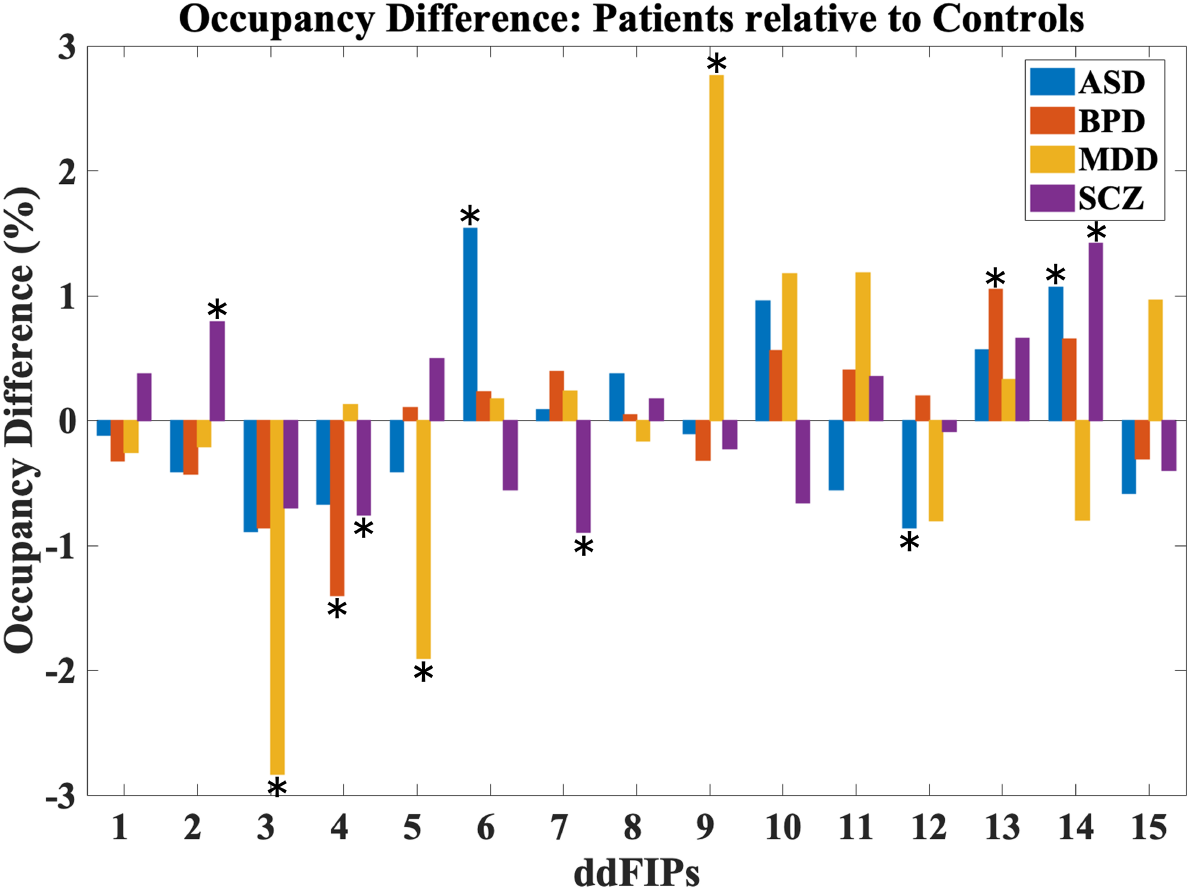
Occupancy analysis results. This figure illustrates the percentage difference in state occupancy across 15 ddFIPs for four clinical groups—ASD, BPD, MDD, and SCZ—compared to HCs. Positive bars indicate increased engagement in a state for the patient group, while negative bars indicate reduced engagement. The significant states relative to controls are shown with an asterisk.

**Figure 4:**
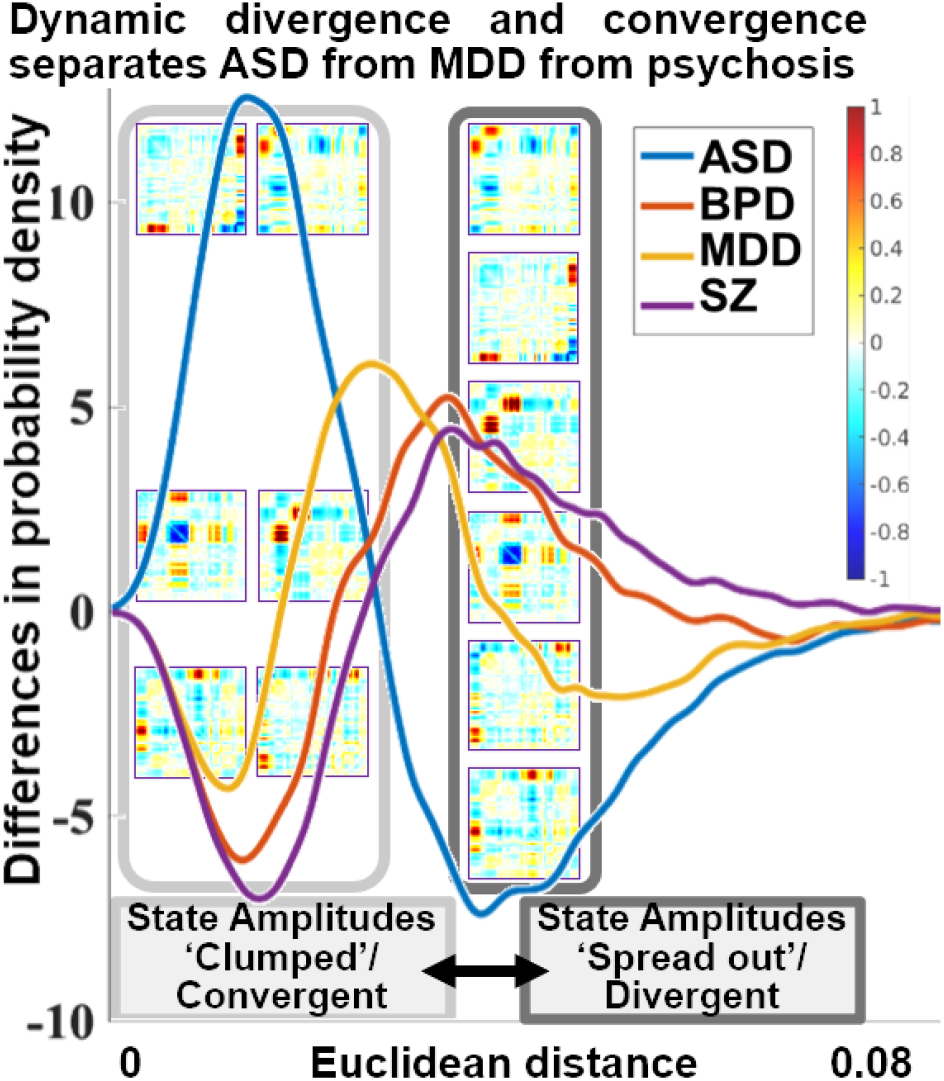
The distribution of pairwise Euclidean distances for SCZ, ASD, BPD, MDD relative to HC, illustrating the variability in connectivity patterns across groups. A wider distribution suggests greater divergence in state amplitudes, indicating more heterogeneous connectivity patterns within a group, whereas a narrower distribution reflects greater convergence and uniformity. This provides a threshold-free visualization of state amplitude divergence, capturing differences in connectivity without applying predefined cutoffs.

Across all groups, some states, such as state 14, show higher average occupancy percentages, suggesting these components are more likely to dominate over time for individuals across all groups. This consistent pattern might indicate core ddFIPs that play a more stable role in brain function across all clinical conditions.

In ASD, significant differences were observed in states 6, 12, and 14. Specifically, the positive occupancy values in these states suggest that individuals with ASD engage more frequently in these connectivity states compared to healthy controls. State 6, for example, involves networks related to sensory processing and attentional focus, aligning with known characteristics of ASD, such as hyper-focus and difficulties with attentional shifts. In state 14, the increased engagement reflects atypical connectivity in networks involved in social and perceptual processing, such as the visual cortex, which are often disrupted in individuals with ASD.

The BPD group exhibits a noticeable peak in state 13, which shows a significantly higher average occupancy percentage than other states in this group. This peak may reflect a tendency toward stronger engagement in specific networks, linked to mood dysregulation and emotional processing associated with BPD. Such heightened occupancy is consistent with the emotional instability and dysregulated mood networks that characterize BPD. The episodic nature of mood fluctuations in BPD may be reflected in this connectivity pattern, supporting the idea that certain connectivity states could serve as biomarkers for mood regulation abnormalities.

For MDD, significant occupancy differences were observed in states 3, 5, and 9. In states 3 and 5, MDD patients exhibit reduced occupancy relative to HC, indicating less frequent engagement in these connectivity states. State 3 corresponds to networks involved in cognitive control and attentional processes, and the reduced engagement aligns with known cognitive and attentional impairments in MDD. Similarly, lower occupancy in state 5 suggests decreased involvement in networks related to reward processing. In contrast, MDD patients show increased occupancy in state 9, which may represent persistent engagement in a state associated with negative self-referential processing including DMN, a common feature of MDD.

Patients with SCZ exhibit significant occupancy differences in states 2, 7, and 14. SCZ patients have lower occupancy rates in state 7, suggesting reduced engagement in connectivity patterns related to cognitive organization and sensory integration, respectively. States 2 and 14, however, show increased occupancy in SCZ, similar to ASD, indicating greater engagement in connectivity patterns linked to self-referential thinking and social cognition, including CC and DMN. This finding aligns with the well-documented social and cognitive impairments in SCZ, particularly in processes related to self and social interaction.

### 3.3 Distribution of Pairwise Distances

To analyze the variability of connectivity patterns across groups, we computed the distribution of pairwise Euclidean distances for each subject to visualize state divergence versus convergence. This provides a threshold-free visualization of divergence and convergence, as it does not rely on the predefined thresholds used in earlier analyses. We generated histograms comparing patients and healthy controls, where the x-axis represents the range of distances and the y-axis shows the probability density. This approach allows us to assess the distribution of connectivity convergence (closeness or similarity of state weights) and divergence (difference in state weights) within each group without applying an explicit cutoff. A wider spread in the histogram indicates greater diversity in connectivity patterns, while a narrower distribution suggests more uniformity in state weights, as shown in Figure 6.

The ASD curve (in blue) illustrates a pronounced peak and a higher density of values in the lower range of Euclidean distances compared to HC. This indicates that individuals with ASD are more likely to exhibit connectivity patterns that are more convergent or similar across time, leading to a ‘clumping together’ of states. The reduced likelihood of values in the higher range of Euclidean distances further supports the idea that brain patterns in ASD are less diverse over time.

For BPD (in orange), the density difference curve demonstrates moderate fluctuations across the range of distances, with peaks indicating both increased convergence and divergence compared to HC. This balanced pattern of both positive and negative values in probability density suggests that BPD patients have connectivity patterns with increased variability, yet not as extreme as ASD. The moderate amplitude of these fluctuations is consistent with the mood variability characteristic of BPD, reflecting intermediate levels of network instability that could contribute to the episodic nature of mood regulation challenges in BPD.

The SCZ curve (in purple) exhibits a broader and smoother spread of positive density values, with moderate peaks extending across a wider range of distances. This distribution indicates that SCZ patients show generally higher variability in connectivity patterns across various network pairs compared to HC, reflected by the positive density values across a broad range of distances. The smooth, widespread pattern is characteristic of the pervasive disruptions in brain connectivity stability often associated with SCZ. This greater variability may align with the disorganized thought processes and cognitive impairments seen in SCZ, where networks fail to maintain consistent communication patterns, contributing to the clinical presentation of the disorder.

The MDD curve (in yellow) has a narrower range of fluctuations with less pronounced peaks, showing mostly positive values in probability density at moderate distances. This narrower spread and positive skew indicate that MDD patients have moderately increased variability in connectivity patterns compared to HC, without the extreme stability or variability observed in ASD. This profile suggests that MDD is associated with subtle disruptions in network connectivity, potentially aligning with its known symptoms of impaired cognitive control. The absence of extreme variability suggests a more stable network compared to ASD and BPD. This method provides insight into the overall network dynamics and can help identify characteristic patterns or anomalies in functional connectivity specific to each clinical group.

### 3.4 Dynamic Convergence/Divergence Analysis

To examine differences in dynamic convergence/divergence between clinical groups and HCs, we conducted cellwise two-sample t-tests on the dynamic convergence/divergence matrices. Furthermore, we applied false discovery rate (FDR) correction to the computed *p*-values with a threshold of 0.05. Each matrix element represents the statistical significance of convergence/divergence differences between specific component pairs, as indicated by −log_10_(*p*) · sign(*t*), where *p* is the *p*-value, and sign(*t*) is the t-value, indicating the direction of the effect. In the resulting matrices, the upper triangular part displays FDR-corrected results, with statistically significant values marked by asterisks, while the lower triangular part shows the corresponding uncorrected results (Figure 5 and Figure 6).

**Figure 5:**
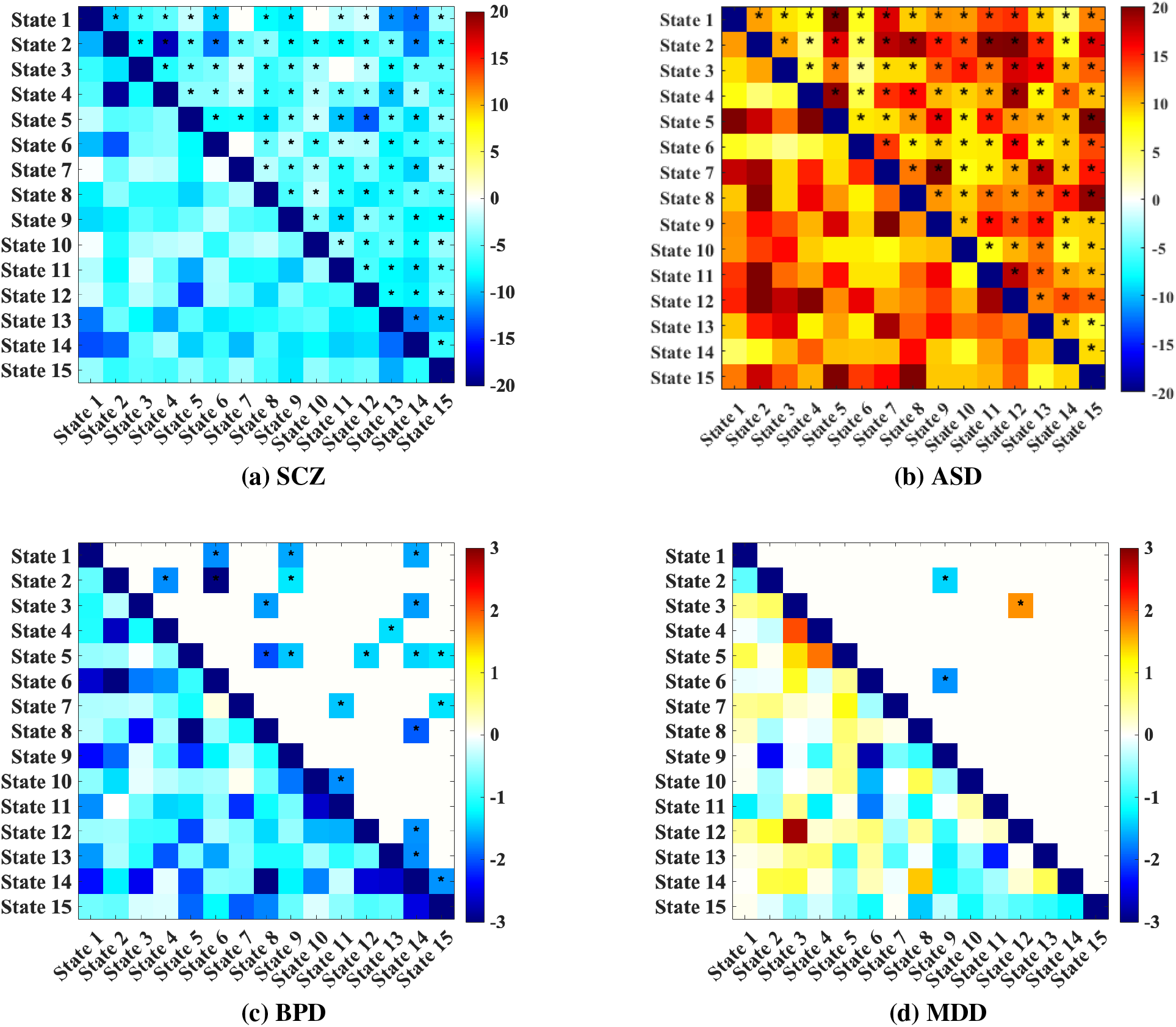
Differences of convergence between a) SCZ, b) ASD, c) BPD, d) MDD and healthy controls. Each matrix element represents the statistical significance of convergence between specific component pairs, as indicated by − log_10_(*p*) · sign(*t*), where *p* is the p-value of the t-test, and sign(*t*) indicates t-value. Significant results after FDR correction are shown in the upper triangular while those significant values before FDR correction are shown in lower triangular part of the matrices. Cells with asterisks denote statistically significant cells after FDR correction.

**Figure 6:**
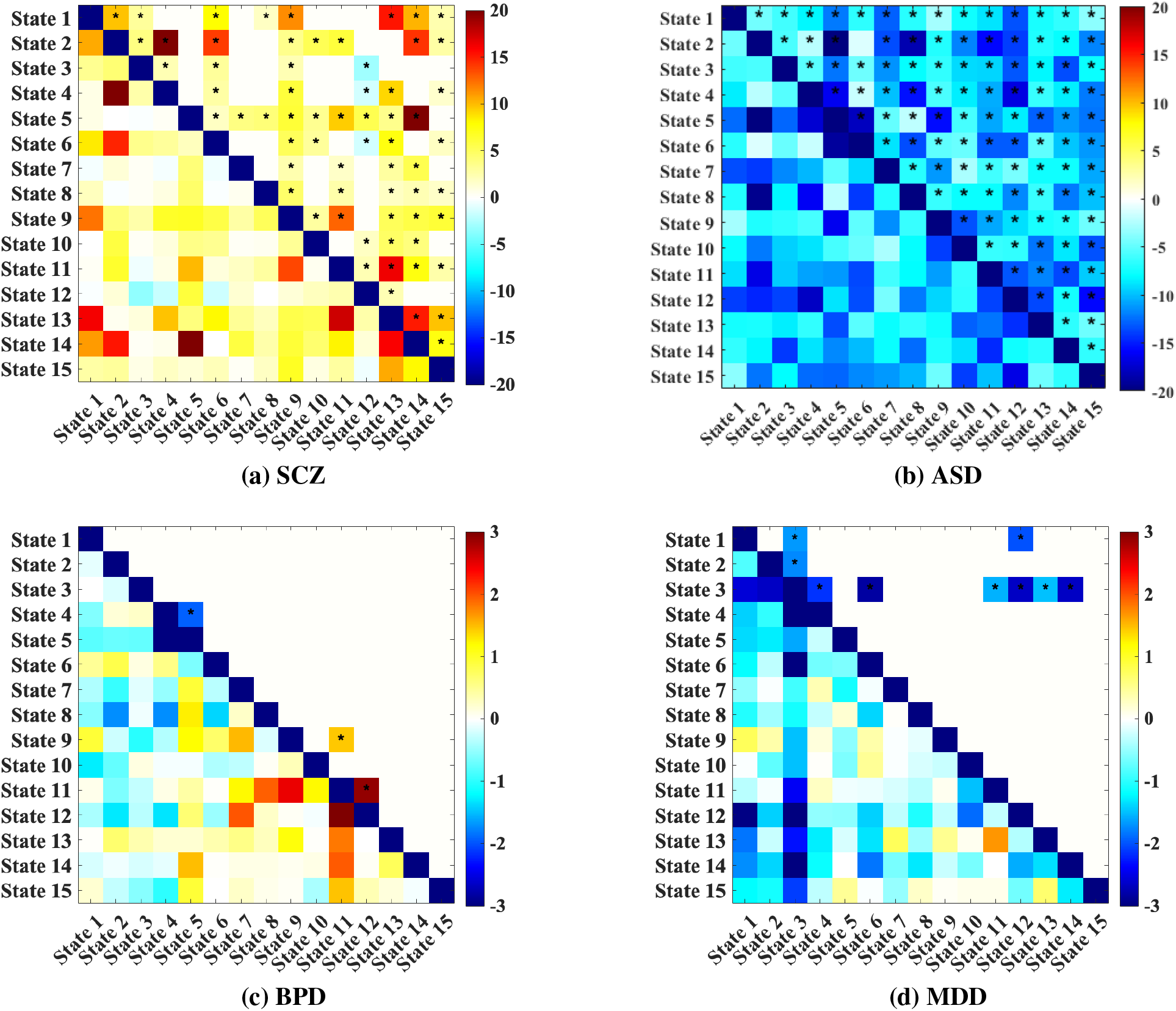
Differences of divergence between a) SCZ, b) ASD, c) BPD, d) MDD and healthy controls. Each matrix element represents the statistical significance of divergence between specific component pairs, as indicated by −log_10_(*p*) · sign(*t*), where *p* is the p-value of the t-test, and sign(*t*) indicates t-value. Significant results after FDR correction are shown in the upper triangular while those significant values before FDR correction are shown in lower triangular part of the matrices. Cells with asterisks denote statistically significant cells after FDR correction.

**Figure 7:**
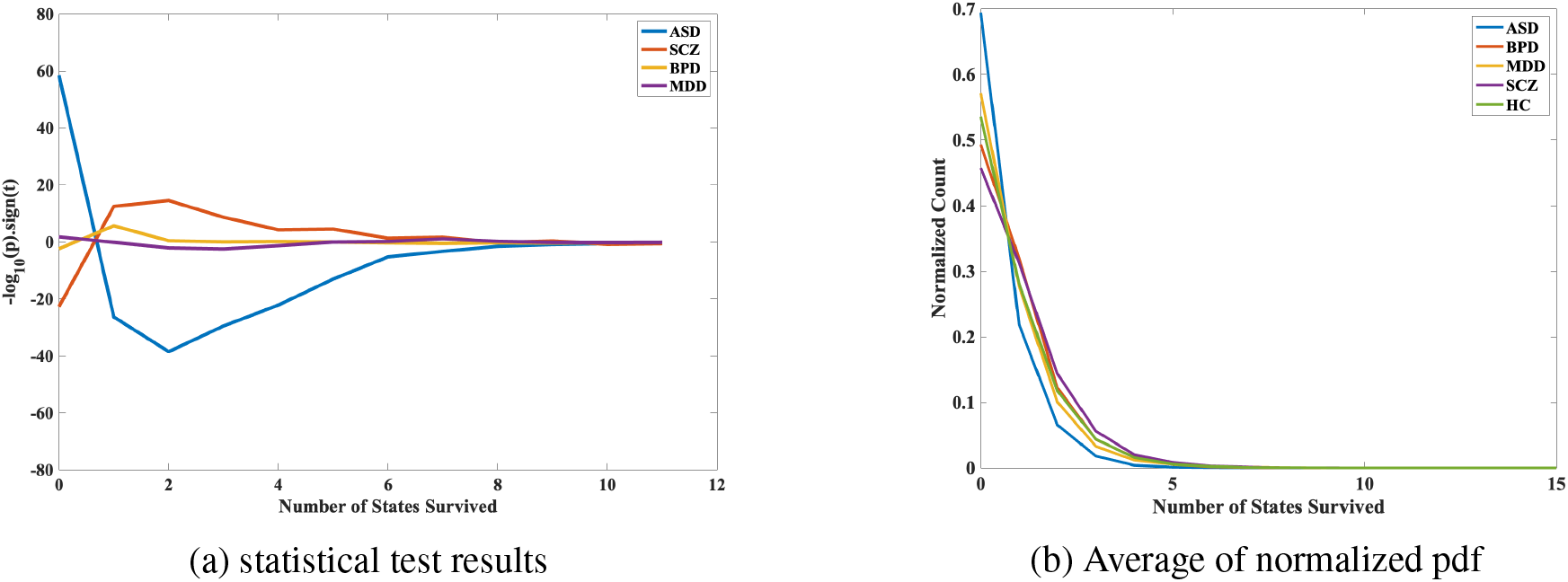
(a) Statistical results represented as − log_10_(*p*) × sign(*t*), illustrating the significance and direction of differences in dynamic state density across various mental disorders. Positive values indicate increased density of states in patients, while negative values indicate decreased density relative to healthy controls. (b) Average normalized probability density functions (PDFs) for state density across mental disorders, showing distinct dynamic connectivity profiles.

**Figure 8:**
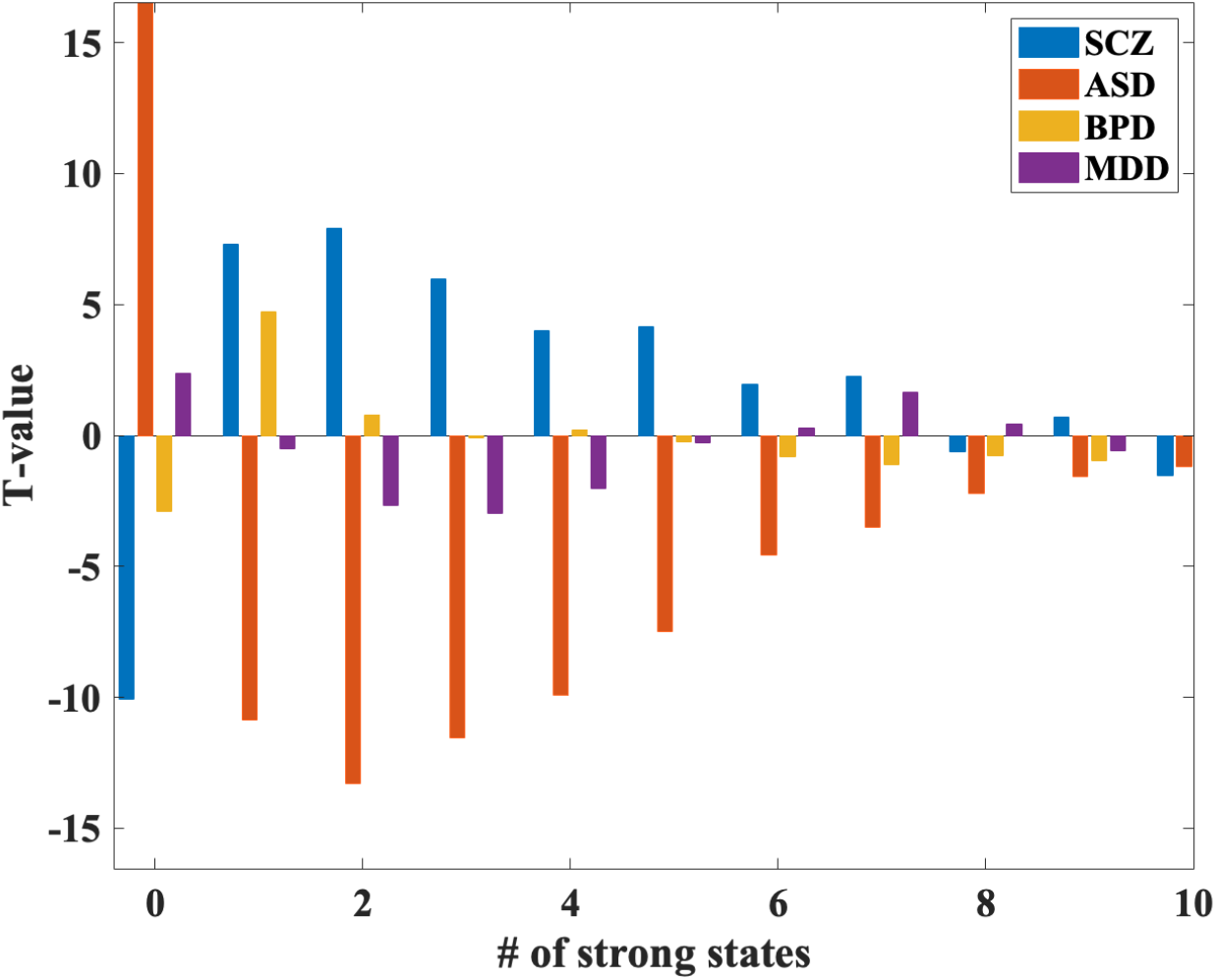
Bar plot showing the distribution of state densities across different mental disorders. SCZ exhibits a high density with fewer instances in the 0-1 states and more in the 3-9 states. BPD shows increased density in the 2-5 states, with fewer instances in the 0 state. MDD demonstrates a decreased density in the 5-6 states, with more instances in the 1-2 states. ASD displays low density, with a higher occurrence in the 0-1 states and lower in the 3-10 states.

#### 3.4.1 Dynamic Convergence Analysis

In the SCZ convergence matrix, significant differences in convergence are observed broadly across state pairs, with mostly negative values. Negative values in the matrix suggest that SCZ patients show lower convergence in certain network pairs compared to HC, suggesting greater variability in dynamic time courses in SCZ. These extensive changes reflect pervasive alterations in connectivity stability characteristic of SCZ. Notably, increased convergence in certain pairs involve frontoparietal and default mode network (DMN) regions, which are frequently implicated in disrupted cognitive control in SCZ (Figure 5 (a)).

The ASD convergence matrix shows more targeted and selective significant differences compared to SCZ. The positive values in several component pairs indicate that ASD patients demonstrate more converged pattern in these pairs relative to HC. The directionality of these effects highlights that ASD patients exhibit strong connectivity in these regions compared to controls, suggesting altered patterns of network interactions (Figure 5 (b)).

The BPD convergence matrix shows fewer but notable areas of significant convergence differences. Specific state pairs exhibit lower convergence in BPD patients compared to HC. These areas of lower convergence may reflect increased variability in connectivity within emotional regulation and limbic networks, which are frequently implicated in mood regulation and dysregulation in BPD (Figure 5 (c)).

The convergence matrix for MDD reveals selective convergence differences with both positive and negative values, indicating varied connectivity stability relative to HC. Positive values in certain component pairs reflect increased convergence in MDD, which may align with enhanced connectivity stability in regions related to negative self-referential processing, such as the DMN. Conversely, negative values indicate lower convergence in MDD patients, potentially reflecting reduced stability in networks associated with cognitive control (Figure 5 (d)).

#### 3.4.2 Dynamic Divergence Analysis

We also evaluated divergence patterns between clinical groups and HC to examine network variability. Dynamic divergence reflects the degree to which timecourses associated with ddFIPs patterns are variable or inconsistent across time within these groups, offering a contrast to the convergence analysis, which focused on stability.

The SCZ divergence matrix (Figure 6 (a)) shows extensive regions with significant divergence differences, indicating that SCZ patients have increased variability across a broad range of component pairs compared to HC.

In ASD (Figure 6 (b)), divergence patterns show relatively significant differences compared to HC, with a marked reduction in variability across component pairs. This reduced divergence in ASD contrasts with the convergence analysis, where ASD showed lower convergence relative to HC. This result suggests that, while ASD patients exhibit less stable connectivity patterns (lower convergence), they do not show excessive fluctuations in connectivity (lower divergence), indicating that the connectivity disruptions in ASD are not driven by variability but rather by atypical consistency.

The BPD divergence matrix (Figure 6 (c)) indicates selective but marked areas of increased variability relative to HC, suggesting that BPD patients experience greater temporal fluctuations in connectivity, particularly within mood regulation networks. This finding is somewhat aligned with the convergence results, where BPD exhibited specific regions of reduced stability, reinforcing the presence of episodic connectivity changes in BPD.

For MDD (Figure 6 (d)), the divergence matrix reveals selective areas of increased variability compared to HC, though less pronounced than in SCZ. The divergence and convergence results present complementary views of connectivity dynamics across disorders.

### 3.5 Dynamic State Density

The analysis revealed distinct patterns of state density across the diagnostic groups, indicating differential dFNC profiles. Individuals with SCZ exhibited a high density of states in mid-to-high state ranges. Specifically, density in states 3 to 9 is significantly more common in this group, while lower-range states (0 to 1) were less frequently occupied. This pattern suggests that SCZ is associated with a shift toward more diverse and dynamic connectivity states, possibly reflecting greater variability in network interactions. In BPD, a marked increase in density was observed, with higher density observed in state counts 2 to 5 compared to other groups. Conversely, zero state was less commonly occupied. This pattern may point to a selective engagement of mid-range connectivity states, highlighting altered but specific dynamic profiles associated with this disorder. Individuals with MDD showed a distinct decrease in overall state density. Density of 1 and 2 were more commonly observed, while state density of 5 and 6 were less common. This suggests a narrower range of dFNC states, reflecting potentially reduced dynamic flexibility in network interactions in MDD. In contrast, ASD was characterized by low density overall, with predominant occupancy in lower-range states (0 and 1). States in the mid-to-high range (3 to 10) were less frequently occupied. This pattern may indicate reduced engagement in diverse connectivity states, highlighting a more rigid and constrained network interaction profile in ASD.

Overall, these results suggest that each diagnostic group is characterized by unique dynamic connectivity profiles, with significant variation in the distribution and density of state occupancies. SCZ and BPD exhibited higher engagement in mid-to-high-range states, reflecting increased variability or specificity, respectively. In contrast, major depressive disorder and autism spectrum disorder showed reduced engagement in these states, suggesting a more constrained and less dynamic connectivity profile.

Figure 9 shows normalized histograms of the distribution of “states survived” for different clusters (cluster 1, cluster 2, and cluster 3) across five groups: HC, SCZ, ASD, BPD, and MDD. Each cluster reflects distinct patterns of state density, highlighting how different mental health conditions vary in their dynamic functional connectivity. Cluster 3 shows a more gradual decline in state density compared to the other clusters, with the shape of the curves varying significantly between patient groups and HCs.

**Figure 9:**
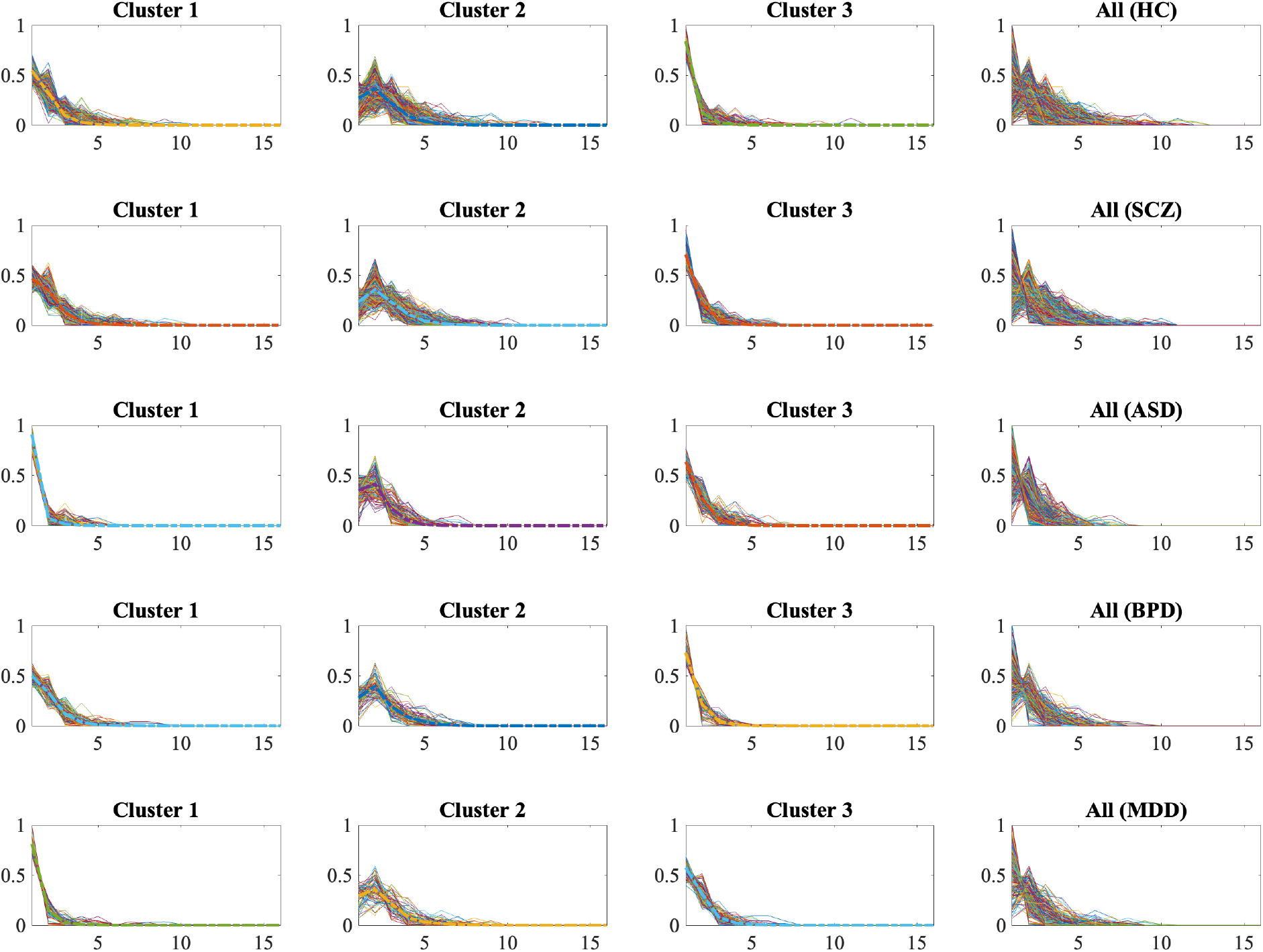
Distribution of “states survived” across three clusters for healthy disorder, BPD: bipolar disorder, MDD: major depressive disorder). Each panel shows the normalized histograms for each cluster, highlighting differences in the dynamic functional connectivity patterns. Notable variability is observed across groups, with HCs displaying more stable distributions compared to the more variable and dispersed patterns in patient groups, particularly for SCZ and BPD. These findings underscore the potential of dynamic state density as a biomarker for differentiating mental health disorders.

Healthy controls tend to have a more uniform and consistent distribution across clusters, particularly for clusters 1 and 3, suggesting a more stable dynamic functional connectivity. The SCZ and BPD groups exhibit more dispersed distributions in all clusters, indicating greater variability and possible dysregulation in state dynamics. The ASD group shows a notably steep decline in Clusters 1 and 2, reflecting fewer sustained dynamic states, which could indicate unique connectivity dynamics. The MDD group displays a slightly flatter distribution in Cluster 2, which may suggest prolonged survival of certain states compared to other groups.

## 4 Discussion

In this study, we introduced and validated a methodological framework, the c-ddFIP, designed explicitly to quantify overlapping and transient dFNC patterns. By systematically applying c-ddFIP to resting-state fMRI datasets from patients with SCZ, ASD, BPD, and MDD, we demonstrated its utility in capturing detailed brain dynamics through metrics such as amplitude convergence, divergence, occupancy, and dynamic state density. The proposed framework effectively differentiated the disorders based on distinctive connectivity dynamic measures, highlighting its potential as a robust analytical tool in neuroimaging research, offering valuable insights into the dynamic functional architecture of psychiatric conditions and laying the groundwork for future efforts aimed at leveraging these metrics for disorder differentiation and diagnostic applications.

SCZ demonstrated the most widespread disruptions, characterized by increased divergence and reduced convergence across a broad range of component pairs. This finding aligns with prior research showing pervasive alterations in connectivity, particularly in DMN and CC networks, which are implicated in self-referential processing, executive function, and adaptive cognition Wang et al. [2015]. The heightened divergence observed in SCZ reflects greater variability in connectivity states over time, suggesting impaired neural stability and adaptability. Such disruptions may underlie hallmark symptoms of SCZ Weber et al. [2020].

Interestingly, our findings also revealed increased occupancy in mid-to-high-range states and selective regions of enhanced convergence. This suggests that while SCZ patients exhibit widespread instability, certain connectivity states may become abnormally stable or over-activated. For instance, hyperconnectivity within DMN and CC network could contribute to the intrusive, self-focused thought patterns observed in SCZ Sasabayashi et al. [2023]. Additionally, our results suggest an imbalance between sensory processing and higher-order networks, with reduced connectivity in sensorimotor regions potentially reflecting impaired integration of external stimuli, a phenomenon previously linked to deficits in perceptual organization Silverstein and Keane [2011].

ASD exhibited a markedly different pattern, with reduced divergence and increased convergence, alongside lower overall state density and occupancy. These findings may reflect atypical neural rigidity, consistent with previous studies reporting reduced dynamic flexibility and atypical stability in connectivity patterns within sensory, attentional, and social networks Rabany et al. [2019]. The predominance of lower-range states and reduced occupancy in higher-order states further highlights the constrained dynamic profile of ASD.

Previous work has identified disruptions in sensory processing and attentional shifts in ASD, particularly involving the visual, auditory, and DMN Lin et al. [2022]. Our findings align with these observations, showing reduced engagement in states involving sensory integration networks. The heightened convergence in specific DMN regions suggests a tendency toward fixed, self-focused cognitive states, which may correspond to behaviors such as repetitive thoughts and difficulty adapting to novel stimuli. Furthermore, atypical stability in connectivity states may explain the rigidity in social and behavioral responses that characterize ASD Hong et al. [2019].

BPD demonstrated increased variability in connectivity, with significant divergence and heightened occupancy in mid-range states. These findings are consistent with the episodic nature of BPD, where mood fluctuations are mirrored by shifts in functional connectivity Cattarinussi et al. [2023]. Previous studies have reported dysregulation in emotional and cognitive networks, including hyperconnectivity within limbic regions and frontoparietal control networks Nguyen et al. [2017]. Our results are consistent with this pattern, showing selective engagement in dynamic states linked to mood regulation.

The increased divergence observed in BPD reflects heightened variability in network interactions, which may contribute to the emotional instability and impulsivity seen in the disorder. At the same time, selective convergence in certain regions may represent compensatory mechanisms aimed at stabilizing cognitive and emotional processes during mood episodes Ishida et al. [2023]. These findings suggest that BPD is characterized by a dynamic interplay between instability and attempts at neural regulation, a hypothesis supported by studies linking mood episodes to shifts in DMN and limbic connectivity Nguyen et al. [2017].

MDD showed reduced dynamic state density and lower occupancy in states associated with cognitive control and reward processing networks. These findings align with prior research documenting hypoactivity in the cognitive control network (CCN) and hyperactivity in the DMN, reflecting a predisposition toward negative self-referential thinking and rumination Gallo et al. [2023]. The increased convergence observed in DMN states suggests persistent engagement in maladaptive thought patterns, a hallmark of depressive symptomatology Albert et al. [2019].

In contrast, the reduced occupancy in mid-to-high-range states reflects diminished engagement in dynamic connectivity patterns associated with cognitive flexibility and reward sensitivity. This may correspond to the anhedonia and executive dysfunction commonly observed in MDD Zheng et al. [2024]. Additionally, the reduced divergence indicates a relative lack of variability in network interactions, suggesting a rigid and inflexible connectivity profile that may limit adaptive responses to environmental changes.

The distinct dynamic profiles observed across disorders underscore the potential of dFNC metrics as biomarkers for psychiatric conditions Du et al. [2017]. Unlike traditional static connectivity analyses, our approach captures transient fluctuations in connectivity, providing a richer understanding of neural dynamics. For instance, the heightened divergence and state density observed in SCZ contrast sharply with the reduced variability and constrained profiles seen in ASD and MDD, highlighting the specificity of these metrics Rabany et al. [2019].

Moreover, the convergence-divergence framework offers novel insights into neural adaptability and resilience. Disorders like SCZ and BPD, characterized by high divergence, may reflect impaired regulation of neural states, contributing to clinical symptoms such as cognitive disorganization or mood instability. In contrast, the increased convergence in ASD and MDD may reflect excessive neural rigidity, limiting the brain’s ability to respond flexibly to changing demands.

These findings also have significant clinical implications. By identifying disorder-specific connectivity patterns, dFNC metrics could aid in differential diagnosis, particularly for conditions with overlapping symptomatology, such as BPD and MDD. Additionally, the ability to quantify dynamic changes offers a promising avenue for tracking treatment responses and disease progression.

Our results align with and extend prior studies emphasizing the importance of dynamic connectivity in psychiatric research Rashid et al. [2014]. The observed disorder-specific patterns reinforce the role of DMN and CCN disruptions as transdiagnostic markers, while highlighting unique dynamic profiles that differentiate disorders. For instance, the reduced divergence in ASD contrasts with the heightened variability in SCZ, suggesting fundamentally different mechanisms underlying connectivity disruptions Rabany et al. [2019].

Future research should explore the longitudinal stability of these metrics, examining how dynamic connectivity changes over the course of illness and in response to treatment. Integrating multimodal data, such as genetic, behavioral, and environmental factors, could further elucidate the complex interplay between brain dynamics and psychiatric symptoms. Additionally, expanding this framework to include other psychiatric and neurological conditions could enhance its generalizability and clinical utility.

## 5 Conclusion

This study introduces and validates a methodological framework to quantify overlapping and transient functional connectivity states derived from resting-state fMRI data. By integrating blind and constrained ICA approaches, ddFIP robustly captures subtle dynamic interactions among intrinsic connectivity networks, overcoming limitations inherent in existing dFNC analysis methods. The proposed metrics, including amplitude convergence, amplitude divergence, and dynamic state density, provide new analytical tools to characterize the brain’s functional flexibility and stability, which are critical aspects often overlooked by static or discrete-state connectivity analyses.

Evaluation of our methodology across large clinical datasets spanning four psychiatric disorders: SCZ, ASD, BPD, and MDD revealed distinct dynamic patterns characterized by the introduced measures: (1) SCZ patients showed widespread divergence, with decreased convergence in sensory networks and increased convergence within higher-order cognitive networks, highlighting disrupted sensory integration alongside excessive cognitive processing states. (2) ASD exhibited significantly increased convergence within and between higher-order cognitive networks, contrasting sharply with reduced convergence (increased divergence) in sensory networks, indicative of atypically stable cognitive states paired with unstable sensory processing. (3) BPD and MDD were characterized by more selective dynamic alterations, showing moderate divergence particularly within emotion-regulation (BPD) and self-referential networks (MDD), indicating targeted, disorder-specific variability in connectivity states. (4) Dynamic state density analysis further clarified these patterns, revealing higher occupancy of diverse connectivity states (greater divergence) in SCZ and BPD, whereas ASD and MDD demonstrated restricted dynamic repertoires with fewer, more convergent states, reflecting differences in neural adaptability and network flexibility.

In summary, this methodological advancement represents a significant step forward in dynamic connectivity analysis, providing robust tools for neuroscience research. Continued development and validation of this and related approaches will be essential for translating dynamic connectivity biomarkers into practical clinical use, thereby advancing the characterization of different mental disorders.

## 6 Author contributions

Conceptualization: NS, VDC; methodology: NS, VDC; software: NS, VDC; validation: NS, SLW, VDC, AI, GP; formal analysis: NS; writing—original draft preparation: NS; writing—review and editing: VDC, SLW, AI, GP; visualization: NS, VDC; supervision: VDC; project administration: VDC; funding acquisition: VDC, AI, GP.

## 9 Acknowledgements

All participants provided informed consent at the time of data collection.

